# Divergent projections from Locus Coeruleus to the cortico-basal ganglia system and ventral tegmental area of the zebra finch

**DOI:** 10.1101/2022.05.20.491849

**Authors:** Jonnathan Singh Alvarado, Jordan Hatfield, Richard Mooney

## Abstract

The locus coeruleus (LC) is a small brainstem nucleus that is the primary source of noradrenaline (NA) throughout the vertebrate brain. Understanding the anatomical organization of the LC is important because NA plays a central role in regulating arousal, attention, and performance. In the mammalian brain, individual LC neurons make highly divergent axonal projections to different brain regions, which are distinguished in part by which NA receptor subtypes they express. Here we sought to determine whether similar organizational features - divergent LC projections acting regionally through different receptor subtypes - characterize cortico-basal ganglia (CBG) circuitry important to birdsong learning and performance. Single and dual retrograde tracer injections reveal that single LC-NA neurons make divergent projections to both cortical and basal ganglia components of this circuit, as well as to dopaminergic nuclei that innervate this circuit. Moreover, in situ hybridization revealed that differential expression of the α_2A_ and α_2C_ adrenoreceptor distinguish LC-recipient song nuclei. Therefore, LC - NA signaling in the songbird CBG circuit could employ a similar strategy as in mammals, which allows a relatively small number of LC neurons to exert widespread yet distinct effects across multiple brain regions.

**Key Points:**

1. The locus coeruleus projects to most of the song system (HVC, RA, LMAN, Area X, DLM).
2. Noradrenergic receptors are regionally specialized in BG despite divergent connectivity of LC neurons.
3. Noradrenergic projections to dopaminergic nuclei could influence vocal variability and learning.

## Introduction

Noradrenergic signaling in the vertebrate brain is an important regulator of arousal, attention, and performance^1,2^. Although noradrenaline (NA) release occurs across the entire brain, the cell bodies of almost all NA-releasing neurons are located in one relatively small pontine nucleus, the locus coeruleus (LC). Thus, one important question is how this small population of NA-releasing neurons exerts a brain-wide influence. Viral-genetic tracing studies in mice revealed that individual LC neurons extend axons that terminate in a large number of different forebrain targets, and this highly divergent architecture is speculated to underlie LC-mediated control of brain states^3^. Another key organizational feature of NA systems in the mouse is the heterogeneous expression patterns of NA receptor subtypes, which allows the LC to exert different effects in different brain regions^4^. Here we sought to explore whether similar organizational features - divergent projections of single LC neurons and heterogeneous NA receptor expression patterns - also characterize how the LC interacts with song-specialized cortico-basal ganglia (CBG) circuitry in the male zebra finch, a small Australian songbird.

Songbirds and zebra finches have proven extremely useful organisms in which to identify neural mechanisms that give rise to the learning and execution of complex vocal behaviors. Male zebra finches learn to sing by copying a tutor song, then use these learned vocalizations as an integral part of a courtship display to attract females. The songs that juveniles sing during the copying phase are characterized by a high level of variability, a feature thought to be important to reinforcement mechanisms of song learning^5^. While song variability declines with the onset of sexual maturity, adult males sing more variable songs when alone but sing more stereotyped songs to females, and this increased stereotypy is more effective as a courtship signal^6,7^. Thus, identifying the neural mechanisms that regulate song variability is important to understanding both juvenile song learning and adult song communication.

Decades of research have delineated a network of brain nuclei that serve a special role in singing and song learning^8^. This song system includes a song motor pathway, which is obligatory for singing, and a song specialized CBG circuit, which comprises the lateral portion of the magnocellular nucleus of the anterior neostriatum (LMAN; a cortical analogue), the thalamic nucleus DLM, and the basal ganglia region Area X. The CBG is essential to song learning and is a major site where song variability is generated and regulated. Notably, NA-mediated suppression of Area X activity is a major means by which the male achieves a highly stereotyped song performance to a nearby female^9^. Here we sought to address whether individual LC neurons make divergent projections to LMAN and Area X, which could provide an efficient means for the LC to coordinately control CBG activity and thus regulate song variability.

Activation of the LC can exert differential effects depending on the brain region. In rodents, for example, LC activation leads to improvements in signal-to-noise in cortical regions^10–12^, increased activity correlations in the striatum^13^, and increased excitability in the basolateral amygdala^14^. These observations raise the question of how the small numbers of LC neurons with highly divergent axonal projections exert disparate effects in different brain regions. One potential explanation is that different populations of LC-recipient neurons express distinct subtypes of NA receptors. Therefore, beyond establishing whether divergent LC connectivity exists in songbirds, we also sought to determine whether LMAN and the Area X express different subtypes of NA receptors. A related issue is whether LC communicates with neuromodulatory systems that also provide input to CBG circuitry. In fact, dopamine (DA) release from ventral tegmental area and substantia nigra pars compacta (VTA/SNc) axons into the Area X is necessary for juvenile song copying which^15^, as previously noted, is a process marked by a high degree of vocal variability^16^. Furthermore, activating NA receptors in the Area X of the adult male suppresses song variability^9^, raising the possibility that these two systems operate in a partially opposing manner. While evidence in mammals indicates that LC activity directly suppresses DA neurons in the VTA^17^, whether the VTA/SNc neurons in the finch receive LC input and express NA receptors both await determination. Thus, we also analyzed the anatomical organization of LC axon terminals and NA receptors in the VTA/SNc complex in the adult male zebra finch, while also characterizing the transmitter phenotypes of both LC neurons and VTA/SNc neurons that receive LC input. These studies indicate that individual LC neurons make divergent projections to LMAN and the Area X, that neurons in these two parts of the CBG circuit express different NA receptor subtypes, and that the LC and the VTA/SNc form a recurrent circuit in which LC is equipped to suppress the activity of DA-releasing neurons in the VTA/SNc.

## Materials & Methods

### Animals

Adult (≥90 days post-hatch) male zebra finches (n = 25) were obtained from the Duke University Medical Center breeding facility. All experimental procedures were in accordance with the NIH guidelines and approved by the Duke University Medical Center Animal Care and Use Committee

### Stereotactic Injections

Male zebra finches were food deprived for 30□min and then anesthetized with 2% isoflurane gas before being placed on top of a small heating pad in a custom stereotaxic apparatus. Rate of breathing and stability of the surgical plane were monitored throughout surgery. The feathers over the skull were trimmed and topical anesthetic (0.25% bupivacaine) was applied before an incision was made in the skin from anterior to posterior with a scalpel. Craniotomies were made with a smaller scalpel at a predetermined distance from the bifurcation of the midsagittal sinus. See Table 1 for stereotactic coordinates relative to the midsagittal sinus. The appropriate depth of anatomical targets was confirmed via electrophysiological recording when available (Differential A-C Amplifier 1700, A-M Systems). To inject tracer, a pressure-based system (Drummond Nanoject II) was used. The injections were either a fluorescent dextran amine or a recombinant cholera toxin subunit B tracer: (Dextran-405; CTB-488, CTB-594, CTB-647, Invitrogen #C22841, #C34777, #C34778). We chose our injection volume based on the size, accessibility, and proximity to other structures. Smaller and more ventral targets such as the LC, VTA, LMAN, and DLM received 25 nl of tracer. Larger targets such as Area-X, RA, and HVC received a total injection volume of 45nl in 15nl increments. Injections were spaced at 5-minute intervals to avoid backflow along the pipette. After these injections, the incision site was closed with tissue adhesive, and the bird was allowed to recover from anesthesia under a heat lamp.

**Table 1.** Stereotaxic coordinates and birds used.

| Target | AP (mm) | ML (mm) | DV (mm) | Beak Angle (°) | Successfully<br>targeted<br>injections |
| --- | --- | --- | --- | --- | --- |
| Area X | 5.1, then 1.8-2.0 | 1.2-1.6 | 2.6-2.9 | 43, then 72 | 4 / 6 |
| DLM | 1.75 | 1.2-1.4 | 4.1-4.4 | 37 | 2 / 6 |
| HVC | 0.4 | 2.5 | 0.15-0.3 | 45 | 3 / 7 |
| LC | -0.3 | 0.9-1.0 | 5.6-5.9 | 28 | 2 / 9 |
| LMAN | 4.2 | 2.0 | 1.7-1.75 | 50 | 3 / 9 |
| RA | -1.8 | 2.4 | 1.5 | 90 | 2 / 4 |
| VTA/SNc | 1.65 | 0.5-1.5 | 6.2 | 37 | 2 / 8 |

### Immunohistochemistry

One week following these tracer injections, birds were deeply anesthetized with an intraperitoneal injection of pentobarbital solution (Euthasol) and then perfused through the heart with 0.025□M phosphate-buffered saline followed by 4% paraformaldehyde. The brain was then removed from the skull and post-fixed in a cryoprotective formalin sucrose solution (30% sucrose in 4% paraformaldehyde) for 48 hours. The next day consecutive sagittal sections of the cryoprotected brain were cut on a freezing microtome. Slides were washed 3 × 5 minutes in PBS plus 0.025% Triton X-100 with gentle agitation. A subset of sections were treated with a primary antibody overnight at 4 °C in PBS with 1% BSA. Sections were washed again in PBS plus 0.025% Triton X-100 3 × 5 minutes. Sections were then reacted with either anti tyrosine-hydroxylase or anti dopamine-beta-hydroxylase secondary antibody (αTH, 1:1,000, Abcam112, αDBH, 1:500, Immunostar #22806) RT (20–25° C) for 1□h at RT in PBS, followed by three 5-minute washes in PBS and mounting onto slides. Sections were imaged with confocal microscope (710 LSM; Zeiss) after coverslipping with Fluoromount-G (SouthernBiotech).

### In-situ Hybridization

Brain sections were processed as described above, but PBS was replaced with RNAse-free PBS(i.e, DEPC-PBS), and a 20% RT paraformaldehyde fixation step was performed before mounting. In situ hybridization was performed using hybridization chain reaction (HCR v3.0, Molecular Instruments). Dissected brain samples were post-fixed overnight in 4% PFA at 4 °C, cryoprotected in a 30% sucrose solution in RNAse-free PBS (i.e., DEPC-PBS) at 4 °C for 48 hours, frozen in Tissue-Tek O.C.T. Compound (Sakura), and stored at −80 °C until sectioning. 80-µm thick coronal floating sections were collected into a sterile 24 well plate in DEPC-PBS, fixed again briefly for 5 min. in 4% PFA, then placed in 70% EtOH in DEPC-PBS overnight. Sections were rinsed in DEPC-PBS, incubated for 45 min in 5% SDS in DEPC-PBS, rinsed and incubated in 2x SSCT, pre-incubated in HCR hybridization buffer at 37 °C, and then placed in HCR hybridization buffer containing RNA probes overnight at 37 °C. The next day, sections were rinsed 4 × 15 minutes at 37 °C in HCR probe wash buffer, twice in 2X SSCT, pre-incubated with HCR amplification buffer, then incubated in HCR amplification buffer containing HCR amplifiers at room temperature for ∼48 hours. On the final day, sections were rinsed twice with 2x SSCT, then mounted on slides and coverslipped with Fluoromount-G (Southern Biotech). After drying, slides were imaged on a Zeiss inverted 710 laser scanning confocal microscope (40x magnification).

### Image Analysis

Confocal z-stacks were obtained with Leica SP8 710 inverted microscopes. Brightness and contrast were adjusted using ImageJ. ImageJ was used for all processing of z-stacks, including z projections, adjustment of brightness and threshold, and changing of look-up tables (e.g., to convert red to magenta). To quantify the number and soma size of neurons, confocal z-stacks were maximum intensity-projected thresholded. The image was then binarized, and a mask for the DBH or TH channel was applied. Labeling in the other channels (CTB/Dextran) was then manually counted within the masked region. For punctate labeling (such as in the case of RA and DLM injection), we used the Puncta Analyzer plug-in for ImageJ v1.29 after background subtraction with a rolling ball radius of 50 and thresholding. Any cell mask with 3 or more of these puncta was considered to be positive for CTB label. For counting enwrapments, the DBH channel was used to first identify the enwrapments. Each z-stack was turned into a maximum projection, and noise was removed with a despeckle filter. After smoothing h an area of >8 μm2 were counted as enwrapments. TH, VGAT, or VGLUT2 signal was then counted only within the enwrapped areas.

## Results

### LC projections to the song system

We first investigated the anatomical connectivity of the zebra finch LC to each nucleus in the CBG circuit (Area X, DLM, LMAN), and to the CBG’s major input and output nuclei in the song motor pathway (HVC (used as a proper name) and RA (robust nucleus of the arcopallium)). We injected small volumes (< 40 nl, see Materials and Methods) of the retrograde tracer cholera toxin subunit B into one of each of these nuclei in different adult male zebra finches (Figure 1). Injections into any one of these nuclei resulted in retrograde label in LC neurons positive for the NA-synthetic enzyme dopamine beta hydroxylase (DBH) (Figure 1a-j); DBH^+^ cells: Area X, 73%, (236/324 DBH^+^ neurons, 6 hemispheres from 3 birds), LMAN, 78%, (215/270 neurons, 5 hemispheres from 3 birds) HVC, 51%, (160/315 neurons, 5 hemispheres from 3 birds) RA, 34%, (82/237 neurons, 3 hemispheres from 2 birds), DLM, 32%, (54/171 neurons, 3 hemispheres from 2 birds). Given our small injection volumes, we consider these results to underestimate the amount of retrogradely labeled LC DBH^+^ neurons, as we only included cases in which the CTB injection was primarily restricted to the target region, even if coverage was partial (see Table 1). These results identify previously undescribed projections from LC to LMAN and DLM, while extending earlier studies in the canary that showed that the LC projects to HVC and RA^18,19^.

**Fig 1.**
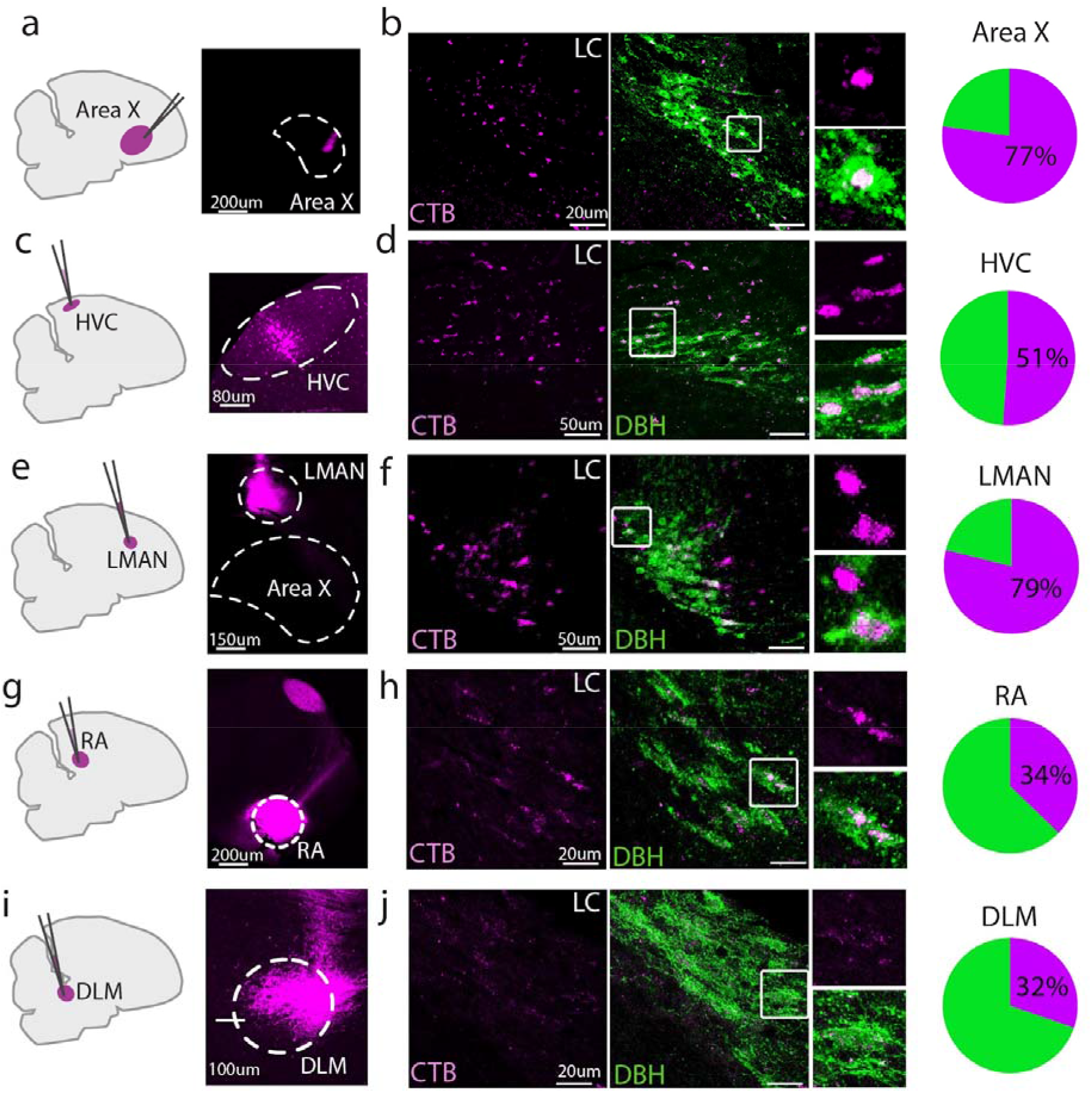
Retrograde tracing from various song nuclei. (a), (c), (e), (g), (i). Sample injection locations into Area X, LMAN, HVC, RA, and DLM. Sagittal section schematic of injection sites. (b), (d), (f), (h), (j). Left, retrograde CTB labeling in LC (magenta), middle, overlay with DBH (green), right, zoomed in view of the boxed region in the right panel, fraction of DBH^+^ cells labeled with CTB for each injected region.

**Fig 2.**
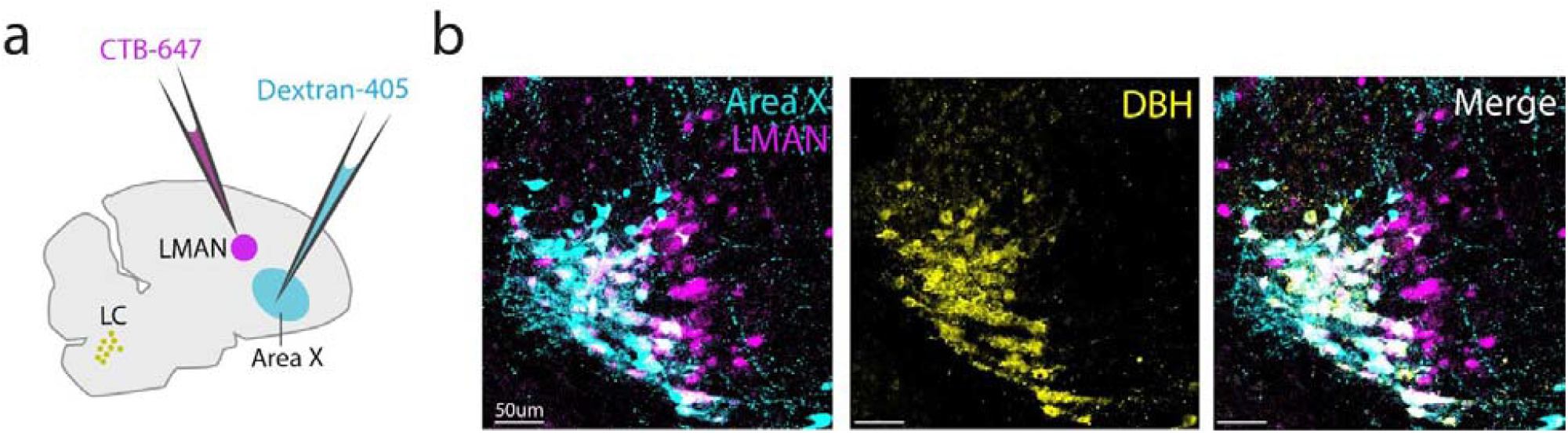
Dual retrograde tracing from LMAN and Area X. (a) Schematic showing CTB-647 (magenta) injection into LMAN, and dextran-405 (cyan) injection into Area X. (b) *Left*: Resulting retrograde label in LC (magenta, LMAN; cyan, Area X; white, double-labeled cells), *Middle*: DBH^+^ cells (yellow) in the same field of view. Right: Merged image, with white denoting triple-labeled cells.

The well-established one-to-many connectivity of LC neurons in mammals led us to ask whether single LC neurons target both Area X and LMAN, which are two major components of the CBG circuit. To answer this question, we injected the fluorescent tracers CTB-647 and dextran-405 into LMAN and Area X respectively and subsequently stained tissue sections containing LC for DBH. We found that 57% of noradrenergic LC neurons (41/89 DBH^+^ neurons, N = 4 hemispheres from 3 birds) were labeled by both tracers. Thus, a substantial fraction of noradrenergic LC neurons make divergent projections to LMAN and Area X.

### α_2A_ and α_2C_ noradrenergic receptor expression in LMAN and Area X

The divergent projections that individual LC neurons make to LMAN and Area X could exert similar or different effects in the regions depending on the expression patterns of NA receptor subtypes^4^. To test this idea, we used in situ hybridization to examine the expression patterns of α_2A_ and α_2C_ adrenergic receptor (α_2A_-AR and α_2C_-AR) subtypes in Area X and LMAN. Notably, this revealed a complementary expression pattern: α_2C_-AR was expressed at high levels in Area X but at relatively low levels in LMAN, whereas α_2A_-AR was expressed at high levels in LMAN but at low levels in Area X (Figure 3a-b). Additionally, both receptors were highly expressed on blood vessels across the brain. Higher power images showed that within Area X, α_2C_-AR was expressed in medium spiny neurons (MSNs), which could be identified by their expression of DARPP-32, while also revealing that a sparse population of DARPP-32 negative cells was enriched for α_2A_-AR (Figure 4a-c). In LMAN, α_2A_-AR was highly expressed in neurons with large somas (median soma diameter of α_2A_-AR^+^ cells, 20.6 ± 3.1 um), which is close to the previously reported size of excitatory LMAN projection neurons ^20^. Taken together, these results indicate that individual LC neurons send divergent projections to LMAN and Area X, and that neurons in these nuclei differentially express α_2A_ and α_2C_ ARs.

**Fig 3.**
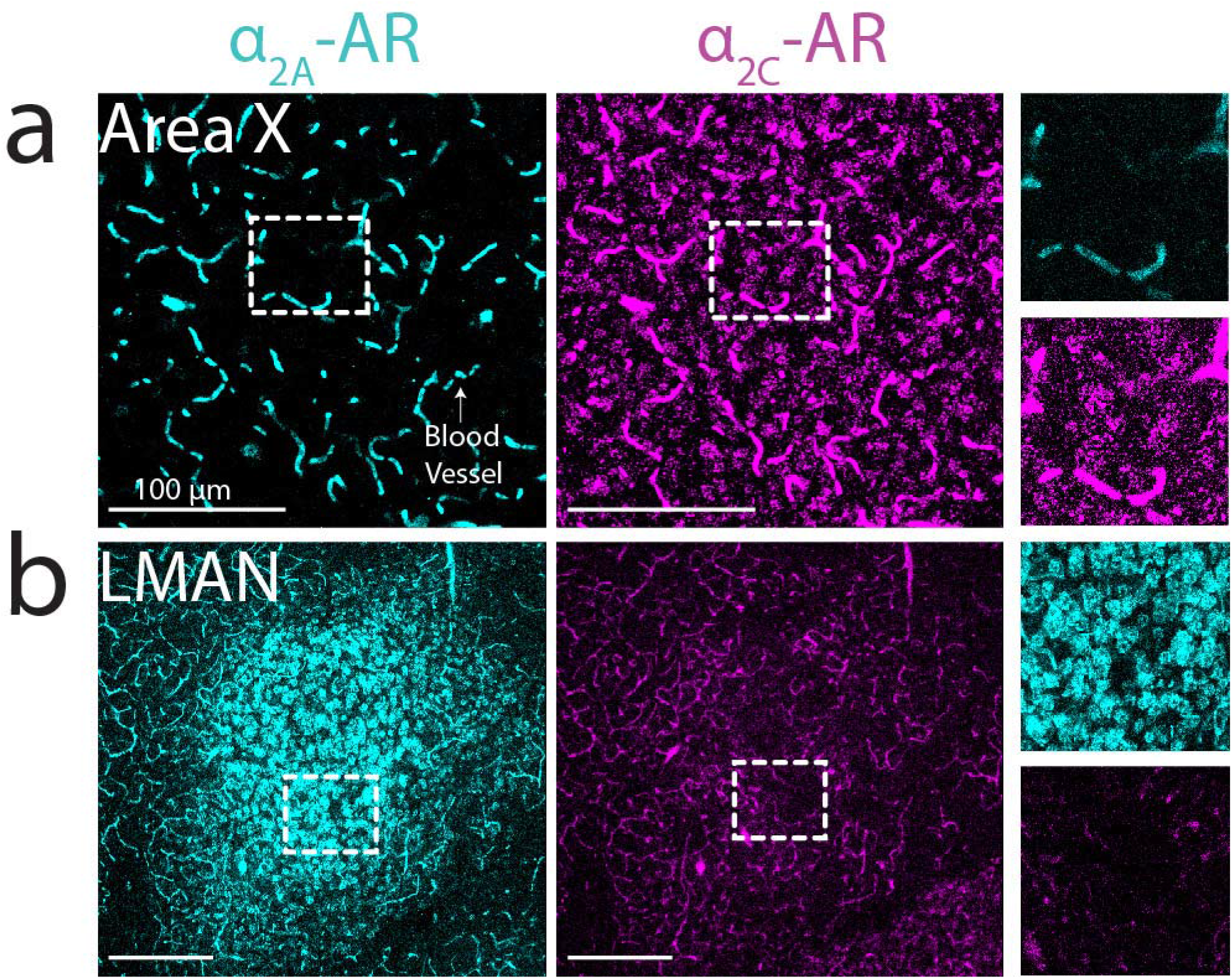
α_2A_-AR and α_2C_-AR mRNA expression in LMAN and Area X. (a) Left: low magnification image of Area X (both receptors were also observed on blood vessels, arrow). Right: zoom-in of white boxed area. (b) Left: low magnification image of LMAN, showing dense expression of _α2A_-AR. Right: zoom-in of white boxed area.

**Fig 4.**
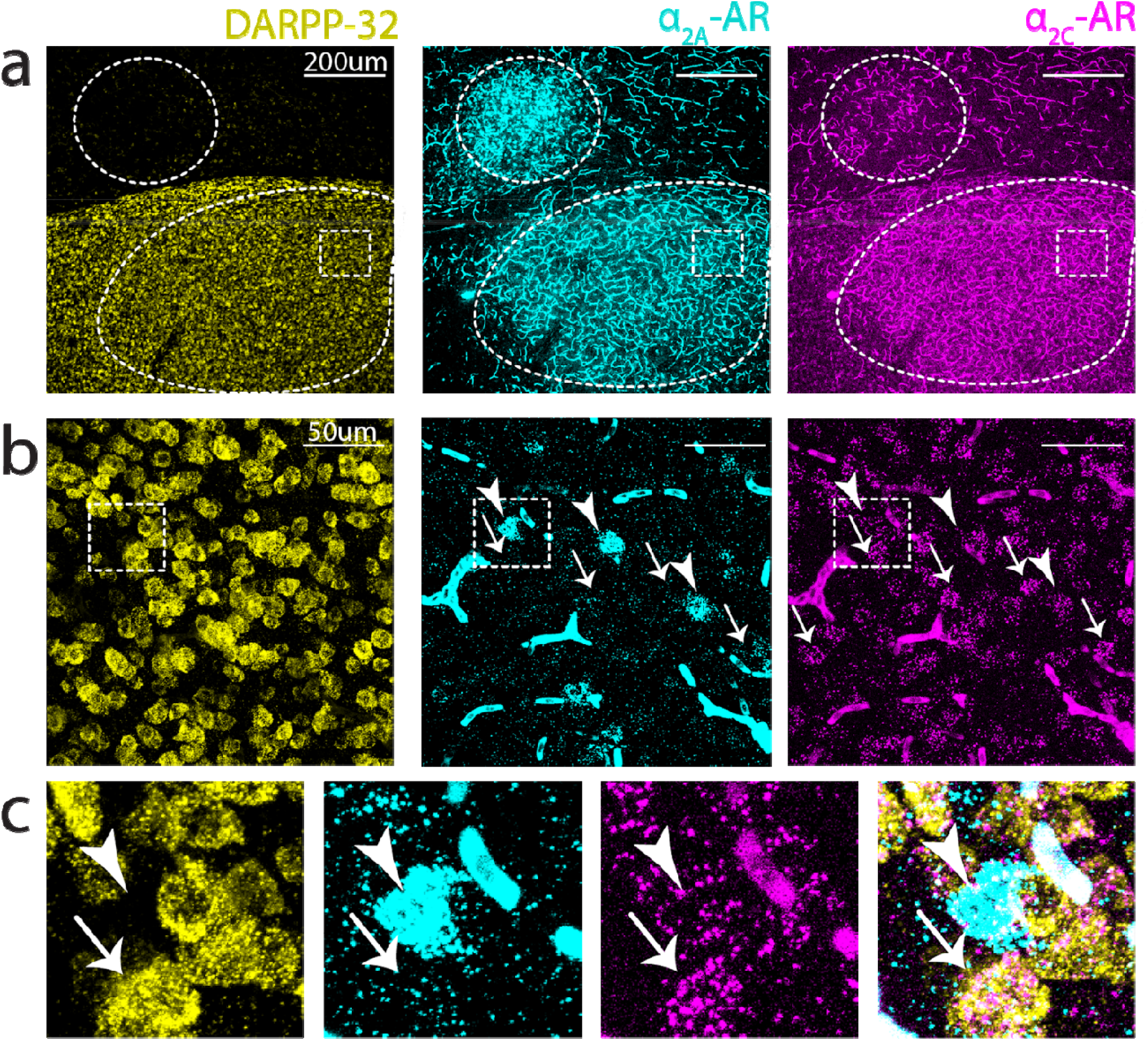
α_2A_-AR and α_2C_-AR mRNA expression within Area X. (a) Low power image showing boundaries of LMAN (top) and Area X (bottom) demarcated by dashed outlines. (b) Higher power image of boxed region in Area X from (a). Sparse clusters of _α2A_-AR can be observed throughout Area X, α_2C_ are more numerous, and show high colocalization with DARPP-32. (c) Zoomed in view of the boxed region in (b) showing that _α2A_^+^ cells do not colocalize with DARPP-32.

### Expression of α_2A_ and α_2C_ adrenergic receptors in VTA/SNc

The VTA/SNc complex is a major source of dopamine (DA) input to both Area X and LMAN. Notably, our in situ hybridization experiments also revealed high expression of α_2C_-AR and α_2A_-AR within the VTA/SNc complex. To better localize the cell types in the VTA/SNc that express ARs, we added a third in situ probe for the enzyme tyrosine hydroxylase (TH), a marker of DA neurons. We found that the majority (160 out of 225, N = 4 hemispheres, 2 birds) of TH+ neurons in the VTA/SNc expressed both α_2A_ and α_2C_ AR mRNA (Figure 5a-b). Additionally, mRNAs for both α_2A_ and α_2C_ ARs were highly expressed in TH+ LC neurons, indicating that ARs could function as autoreceptors to regulate LC activity (Figure 5c). Therefore, NA release from LC neurons could directly modulate the activity of DA neurons in the VTA/SNc.

**Fig 5.**
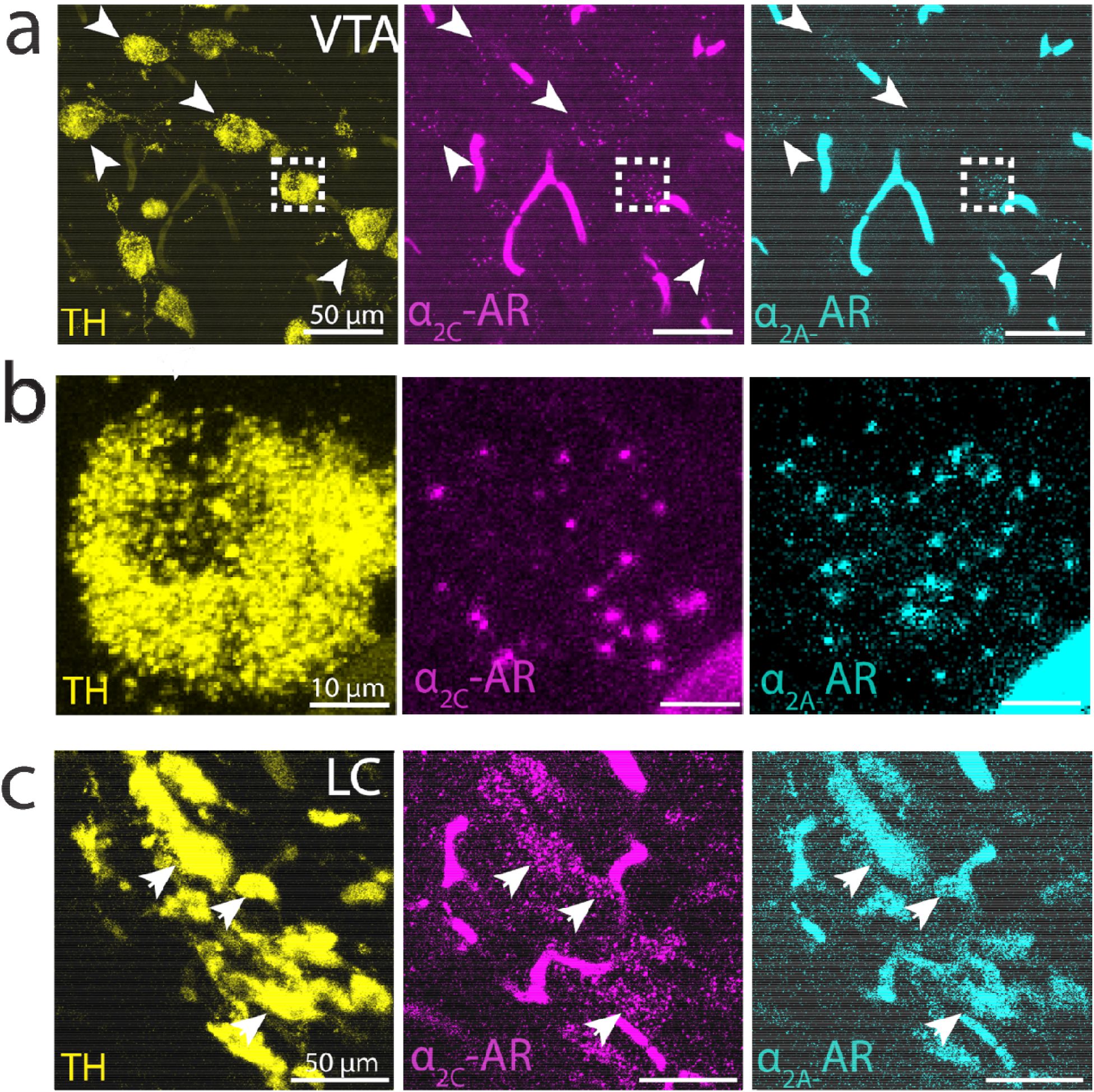
α_2A_-AR and α_2C_-AR mRNA expression in VTA and LC neurons. (a) TH^+^ neurons in VTA (yellow, left) and expression of _α2A_(magenta, middle) and _α2C_ (cyan, right). (b) Magnification of a single cell in boxed region. Arrowheads indicate location of sample TH^+^ cell bodies across images. (c) _α2A_ and _α2C_ receptor expression in LC cell bodies.

### Anatomical projections between LC and VTA/SNc

The distinct patterns of NA receptor expression that we detected in the VTA/SNc suggest that NA release from LC could modulate VTA/SNc activity. Moreover, VTA/SNc DA neurons project to the CBG^21^, indicating that LC could influence Area X activity directly through NA receptors within Area X, as well as indirectly by influencing DA release from VTA/SNc onto Area X. We therefore tested whether LC-NA neurons also project to the VTA/SNc. First, we stereotaxically injected the retrograde tracer cholera-toxin B (CTB) into the VTA/SNc complex of adult male zebra finches. Tracer injections that were largely restricted to the VTA/SNc resulted in retrograde labeling in the majority of DBH^+^ LC cell bodies (80 out of 98 cells, N = 4, hemispheres, 2 birds, Figure 6a-c). Additionally, we consistently observed a strip of retrogradely labeled DBH^−^ neurons dorsal and rostral to the double-labeled cells in LC (Figure 6c.). Thus, the VTA/SNc receives input from DBH^+^ cells in the LC as well as from DBH^−^ cells that surround these noradrenergic LC neurons.

**Fig 6.**
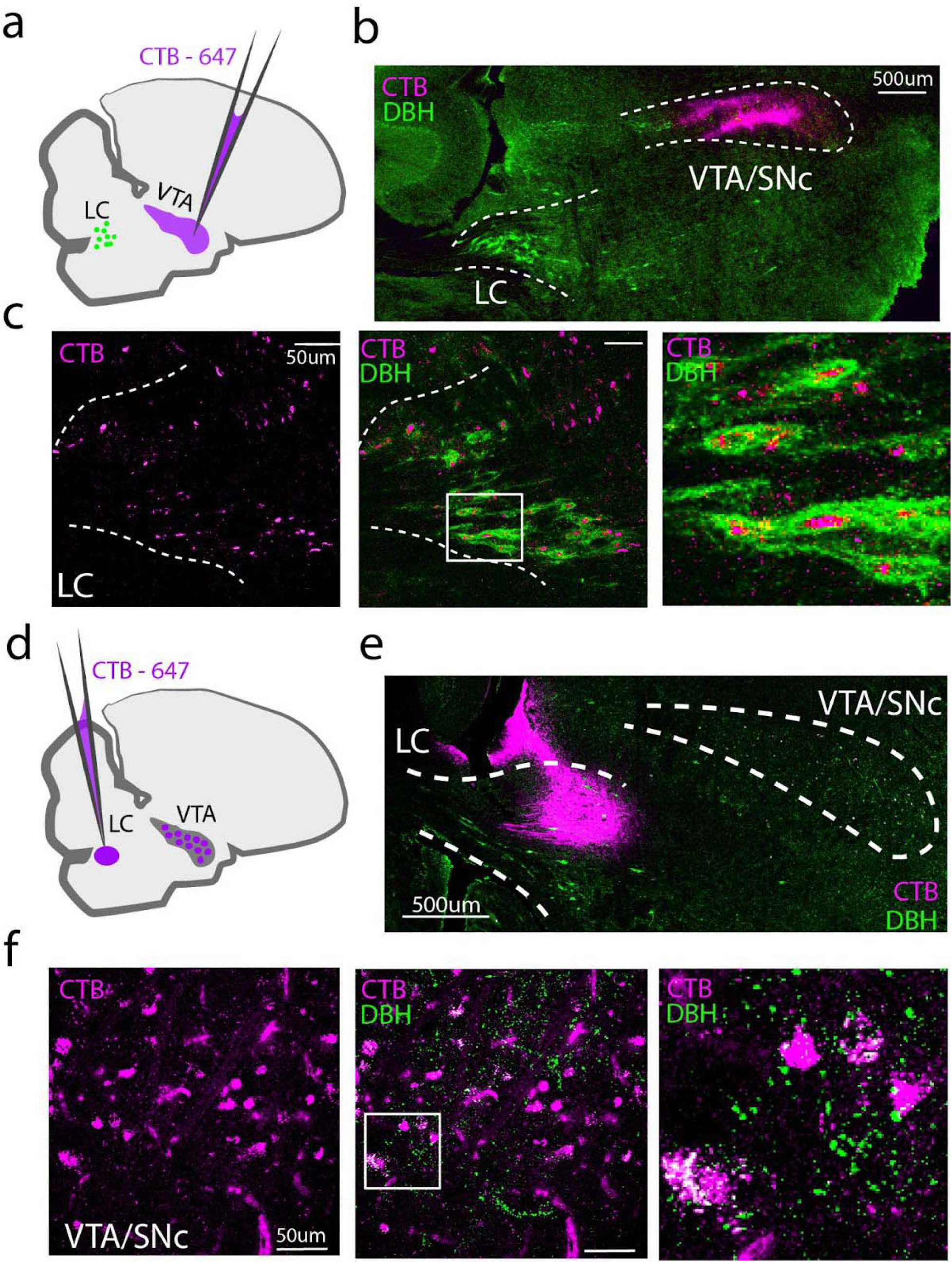
Projections between LC and VTA/SNc. (a) Experimental approach for retrograde labeling of DBH^+^ LC neurons that provide input to the VTA/SNc. A small (10-20nl) injection of CTB was targeted to VTA/SNc. (b) Low power image showing CTB injection in the VTA/SNc. (c) Left: Retrogradely labeled cell bodies (magenta) in LC, middle: merge with DBH stain. Right: Zoomed in view of the boxed region in the middle panel. (d) Experimental approach for retrograde labeling of VTA/SNc cells that provide input to the LC. (e) Low power image showing CTB injection in LC. (f) Retrogradely labeled cells in VTA/SNc. Left: Retrogradely labeled cell bodies from the CTB injected in LC. Middle: merge with DBH^+^ fibers in VTA/SNc. Right: Zoomed in view of the boxed region in the middle panel.

We next tested whether the VTA projects to the LC in the zebra finch, as has been shown in mammals^22^. To avoid lowering our injection pipette through the VTA, we approached the LC through the cerebellum, crossing through the fourth ventricle (See Table 1 and Fig 6d). For birds in which CTB was predominantly limited to the DBH^+^ boundaries of the LC (N = 2 out of 9), we observed retrogradely labeled cells in the VTA/SNc that overlapped with DBH^+^ fibers (Figure 6d-f.) Taken together, these retrograde tracing experiments indicate that the VTA/SNc and LC make reciprocal connections.

### A subset of LC cells that project to the VTA also project to Area X

We next investigated whether individual LC neurons make divergent projections to Area X and the VTA/SNc. Tracer injections of Dextran-405 made in the Area X and CTB in the VTA/SNc produced co-labeled a majority of DBH^+^ neurons in the LC (80/116 DBH^+^ cells were double labeled, N = 4 hemispheres, 2 birds, Figure 7a-c). These results indicate that a substantial number of noradrenergic LC neurons project to both the VTA/SNc and Area X.

**Fig 7.**
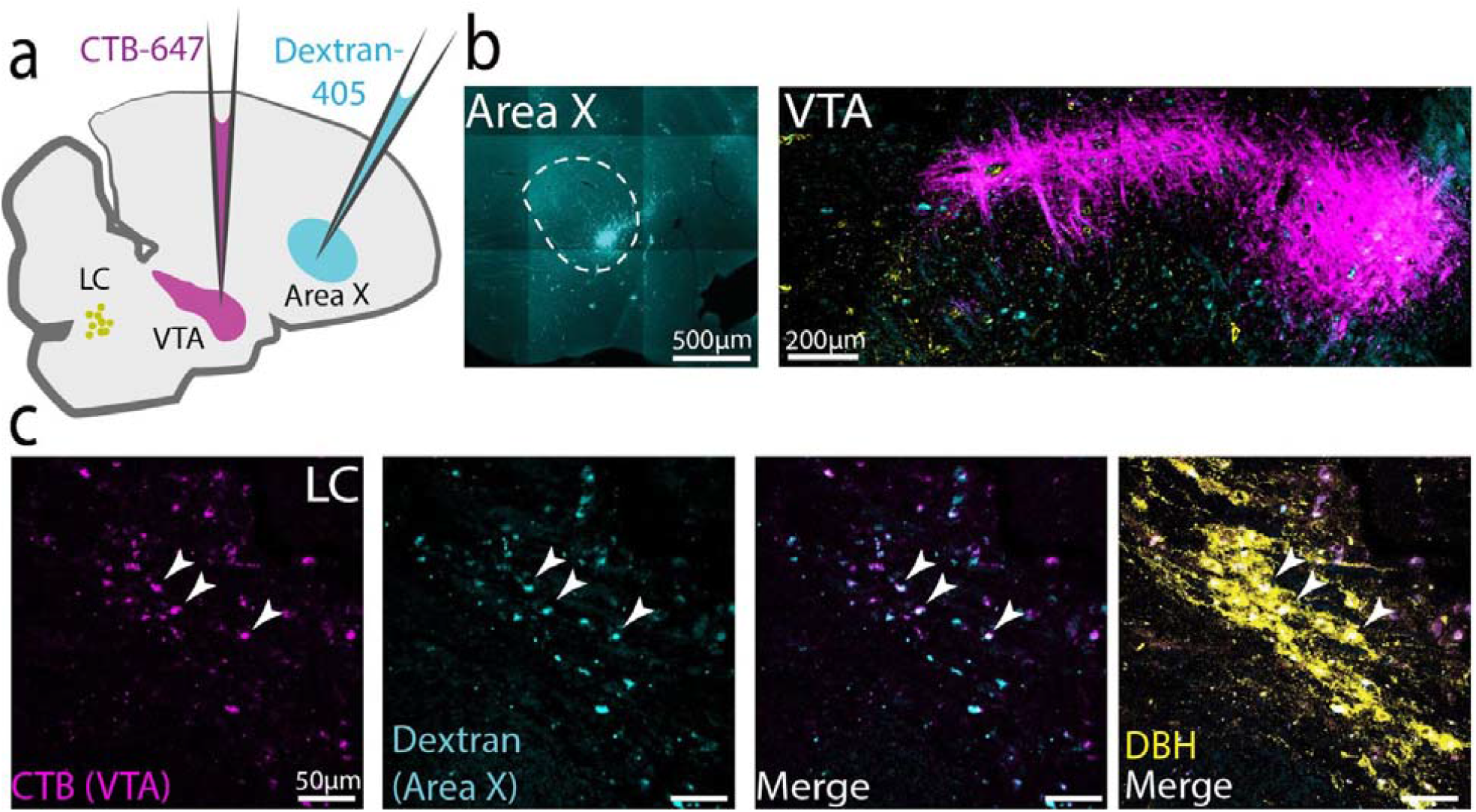
Individual LC NA neurons project to both VTA/SNc and Area X. (a) Experimental approach for dual labeling from Area X (Dextran-405, cyan) and VTA/SNc (CTB-647, magenta). LC cell bodies depicted in yellow. (b) Injection sites in Area X and VTA/SNc (c) High power image of retrograde labeling in LC. From left to right: CTB-647 from VTA/SNc, Dextran-405 from Area X, merge, and overlay with DBH.

### Noradrenergic fibers in VTA/SNc preferentially form appositions with VGAT^+^ cells

Upon closer investigation, we also noticed that DBH^+^ axons formed dense appositions, or enwrapments, around cell bodies within the VTA/SNc complex, especially around TH^−^ cell bodies (Figure 9a-b). To better characterize which cells in the VTA/SNc complex were targeted by these DBH^+^ enwrapments, we performed in situ hybdridization for VGAT and VGLUT2 followed by immunohistochemistry to label DBH^+^ axons. We found that 78% of DBH^+^ enwrapments within VTA/SNc targeted VGAT^+^ cell bodies, but only rarely targeted TH^+^ or VGLUT2^+^ neurons (32/42 enwrapments onto VGAT^+^ cells, 4/42 onto VGLUT^+^ cells, n = 4 hemispheres, 2 birds) (Figure 9c,d). Moreover, only 14% (3/21) enwrapments targeted TH^+^ neurons (n = 4 hemispheres, 2 birds. (Figure 8a,b). These results suggest that LC axons make especially strong synaptic connections with GABAergic cells of the VTA/SNc complex, which prior studies have shown inhibit the activity of DA-releasing VTA/SNc neurons.

**Fig 8.**
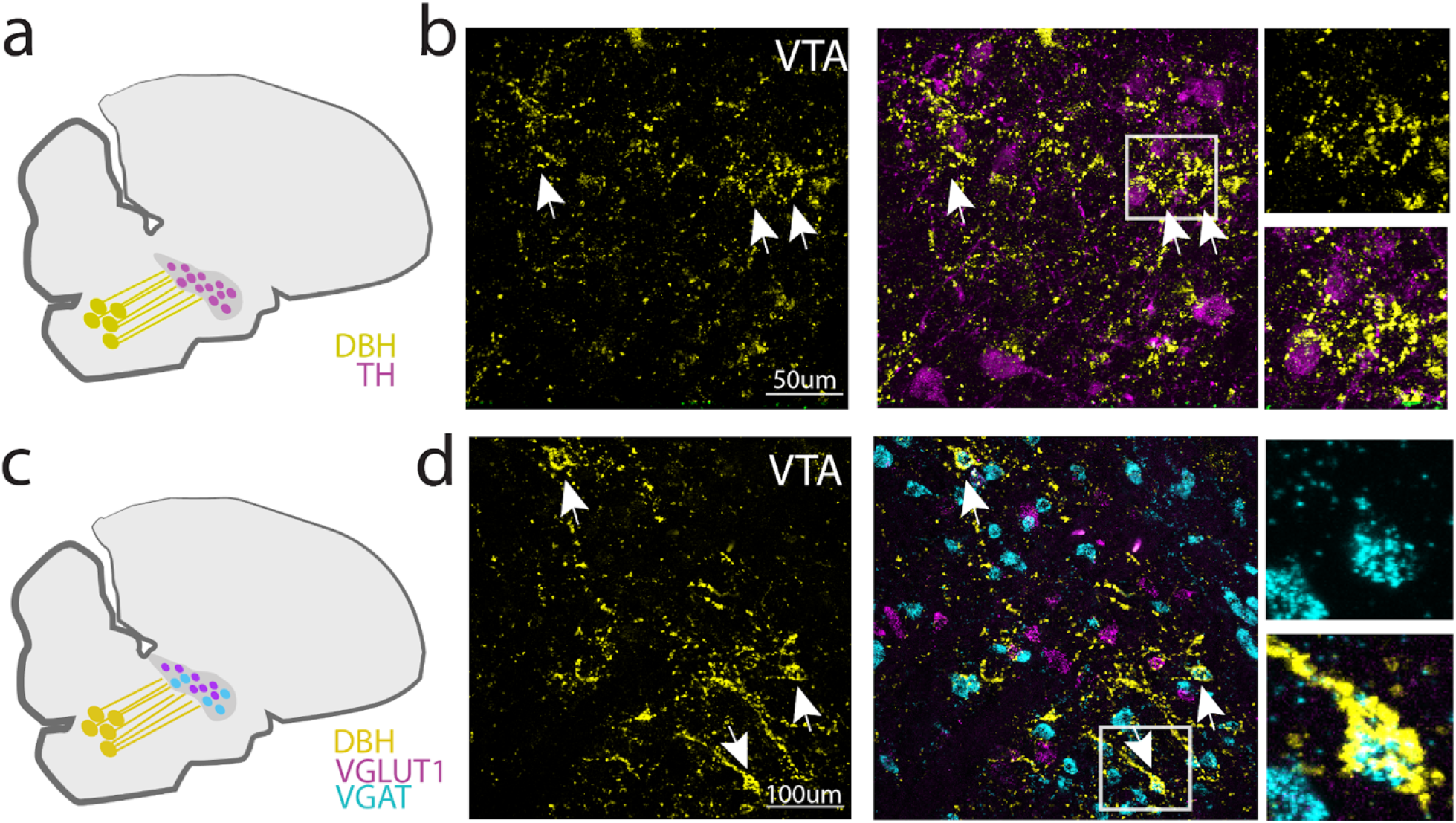
DBH^+^ terminals form perisomatic appositions onto VGAT^+^ neurons in VTA/SNc.(a) DBH^+^ terminals in VTA/SNc were visualized in combination with TH^+^ neurons in VTA/SNc. (b) Left: High power images of VTA/SNc showing DBH terminals (yellow) and enwrapments (arrowheads). Middle: DBH^+^ terminals and TH^+^ VTA/SNc cell bodies. Right: Magnification of boxed region showing TH^−^ enwrapments. (c) DBH^+^ terminals in VTA/SNc were visualized in combination with VGAT and VGLUT2 in situ labeling. (d) Left: High power images of VTA/SNc showing DBH terminals (green) and enwrapments (arrowheads). Middle: DBH^+^ terminals and in situ labeling. Right: Magnification of boxed region showing enwrapments around VGAT^+^ cells.

**Fig 9.**
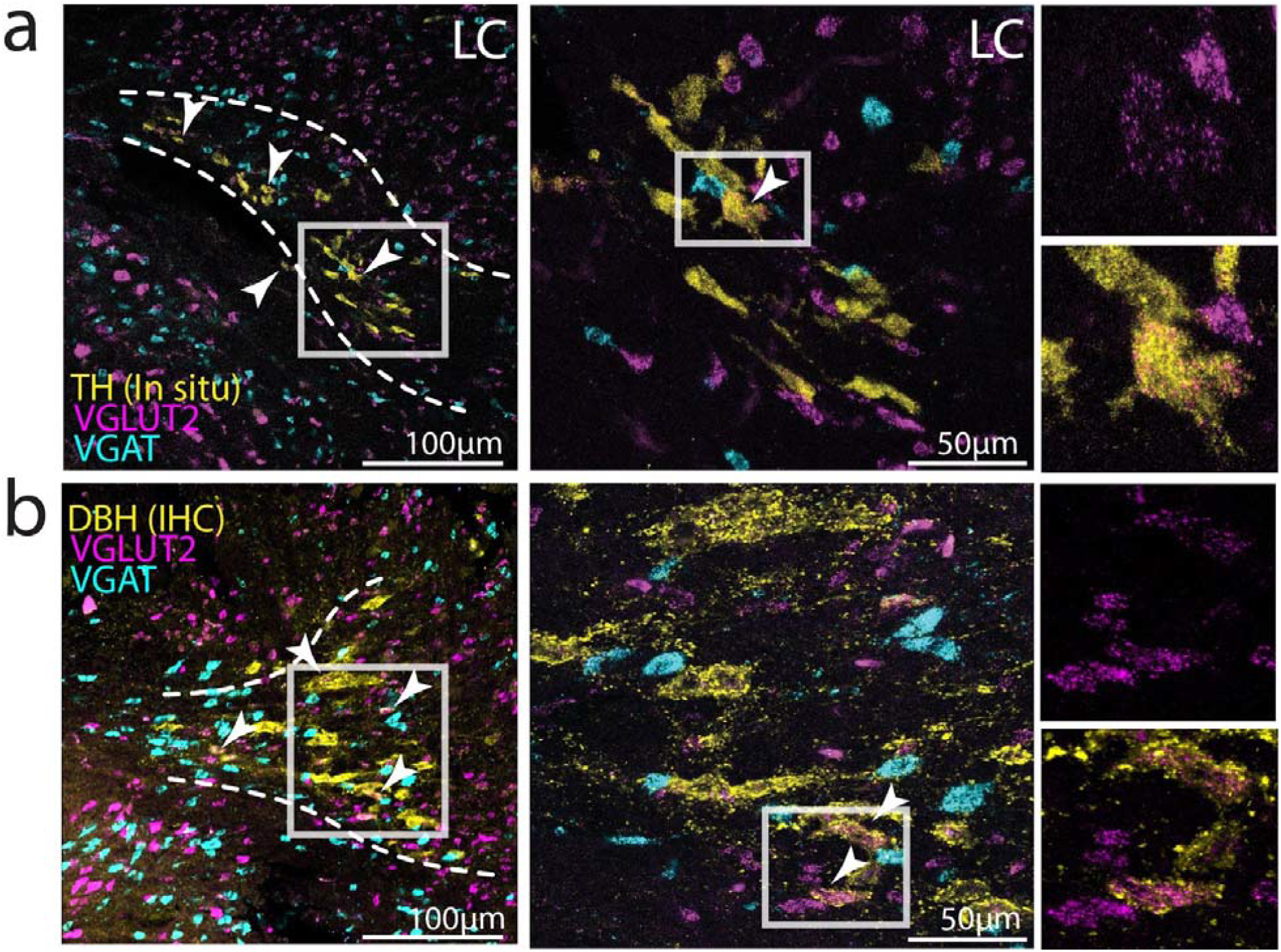
VGLUT2 mRNA expression on LC DBH^+^ neurons. a) Left: Low power image of in situ labeling for TH (yellow), VGLUT2 (magenta) and VGAT (cyan) mRNA expression in LC. Middle: Higher power image of boxed region in left column. Right: Zoom in on example LC-NA cells that co-express VGLUT2. b) Same as in (a), but using DBH protein (yellow) as a marker for LC-NA neurons.

### A subset of LC neurons co-express VGLUT2

Here, we have focused on the expression of NA receptors in the CBG and VTA/SNc complex, but an emerging body of evidence indicates that catecholaminergic neurons can co-release other neurotransmitters such as glutamate and GABA^23,24^. To investigate whether LC neurons in the finch could also release these neurotransmitters, we performed in situ hybridization for the vesicular glutamate and GABA transporters VGLUT2 and VGAT in tissue sections containing LC NA neurons, which we identified using a third in situ probe for TH. We observed that approximately 30% (28/96 neurons, N = 4 hemispheres, 2 birds) of TH+ LC neurons expressed VGLUT2, but none (0/96 neurons) expressed VGAT. We further confirmed this result using a DBH antibody to identify noradrenergic LC neurons (Figure 9b). Therefore, LC neurons in the finch have the potential to release glutamate as well as NA on their postsynaptic targets.

## Discussion

### LC projections targets in the zebra finch

Despite its small size (∼1500 neurons in mice, ∼800 in zebra finches), the LC is known to influence fundamental aspects of behavior, such as wakefulness and attention. One way the LC achieves such influence is through highly divergent axonal projections, an anatomical feature that enables individual LC neurons to communicate with a wide array of brain regions. In support of this idea, recent studies have used input- and output-specific viral tracing to demonstrate that LC neurons receive inputs from many brain regions and send axonal projections to many destinations ^3^. Here we show a similar anatomical organization in the songbird for LC’s outputs (summarized in Figure 10a) and different components of song specialized CBG circuitry, whereby individual LC neurons make divergent projections to Area X, LMAN and the VTA/SNc complex. Therefore, individual LC neurons broadcast their signals to midbrain, basal ganglia, and cortical regions alike in the finch.

**Fig 10.**
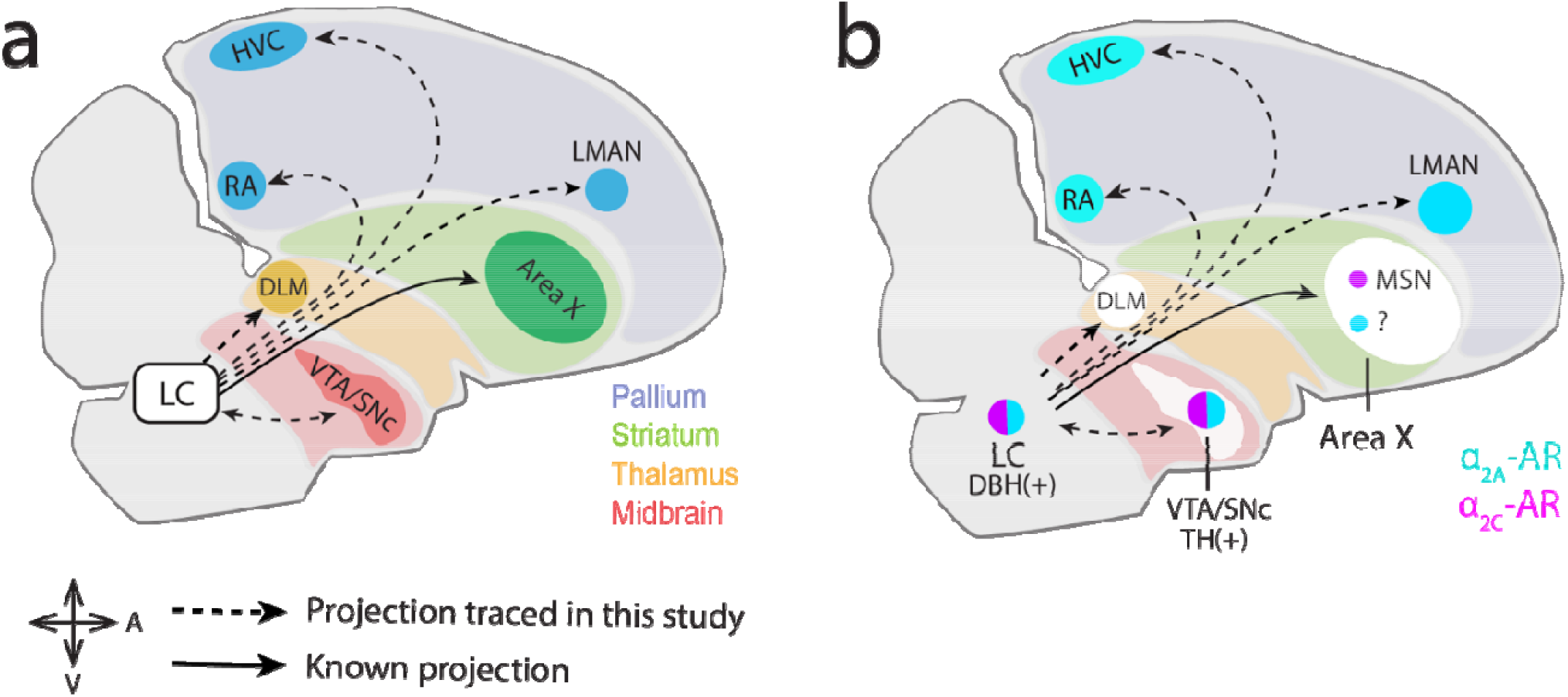
Summary of findings for this study. (a) Parasagittal diagram of the male zebra finch brain highlighting the projections observed in this study. (b) Summary of α_2C_ and α_2A_ receptor expression in the investigated song nuclei.

In addition, we also found that retrograde tracing from another 3 song nuclei (RA, HVC, DLM) resulted in labeling that was largely restricted to LC DBH^+^ cells. These results stand in contrast to previous tracing studies in canaries, which found very sparse LC projections to HVC and RA ^18,19^. More broadly, our results show that LC’s divergent connectivity extends beyond the CBG, to encompass midbrain, thalamic, and pallial regions.

### Differential α_2A_ and α_2C_ AR expression in the CBG circuit

Our in situ hybridization experiments further demonstrated dense expression of the Gi-coupled adrenergic receptors α_2A_ and α_2C_ in LMAN and Area X, respectively, suggesting that NA release onto these areas could act synergistically to inhibit CBG output. Within Area X, NA-mediated activation of α_2C_-ARs directly suppresses SN activity, presumably leading to reduced pause variability in downstream pallidal neurons ^25,26^. In addition, NA application in LMAN leads to suppressed excitatory transmission from LMAN to the motor output nucleus RA (Sizemore & Perkel, 2008), presumably via activation of α_2A_-ARs. Thus, NA signaling could work in parallel across LMAN, Area X and RA to reduce vocal variability during directed song.

### Projections between VTA/SNc and LC

Anatomical tracing revealed mutual connectivity between LC and VTA/SNc, and dual tracing experiments further showed that individual VTA/SNc-projecting LC neurons also project to Area X. We identified two potential ways in which LC activity could modulate VTA/SNc activity. First, noradrenergic terminals within VTA/SNc preferentially formed enwrapments onto VGAT^+^ neurons, potentially allowing the LC to modulate activity of VTA/SNc interneurons, including perhaps through the co-release of glutamate. Second, TH^+^ VTA/SNc neurons co-expressed the two Gi-coupled adrenergic receptors α_2A_ and α_2C_ (Figure 10b). Therefore, the LC could suppress the activity of VTA/SNc DA neurons both indirectly through local interneurons and directly through inhibitory AR receptors.

### Functional and behavioral implications of LC connectivity

The widespread connectivity pattern of LC neurons we observed here, which had not been previously described in songbirds, is well suited to “switch” the song system into different states depending on behavioral context, motivational factors and endocrine cues. For example, zebra finches change their song properties or body movements depending on whether they are practicing, performing ^27,28^, tutoring ^29^, or being tutored ^30^, and many of these behavioral changes are associated with changes in neural activity and immediate early gene (IEG) expression in song CBG nuclei ^26,31,32^, as well as in the LC and VTA/SNc complex ^29,33,34^. In support of a role for LC in facilitating brain-wide changes that control context-dependent changes to singing and other related behaviors, systemic noradrenergic lesions affect courtship behaviors and their corresponding neural signatures ^35,36^, and LC stimulation affects social context-sensitive song features _37_.

### Relevance to courtship singing and song variability

The songs of male zebra finches are faster, less variable and more vigorous when they are directed to a female as compared to when they are undirected (produced in isolation, e.g.) ^27,28^. These acoustic changes are readily detected by female receivers, who strongly prefer directed songs over undirected songs ^6^. Moreover, changes in song production during directed singing are thought to be caused by changes in neural activity, particularly within the CBG. Previous evidence suggests that D1 receptors play a role in this context-dependent switch ^38,39^, and DA levels are slightly higher during directed song than during solo song ^40^. Despite the initial establishment of the DA system as a primary focus of interest for understanding context-dependent changes in song, more recently the LC-NA system has also been implicated in this process. Systemic lesions of the noradrenergic system largely abolish context-dependent differences in immediate early gene (IEG) expression in Area X^35^). Furthermore, pharmacologic activation of noradrenergic receptors can also selectively suppress glutamate release from LMAN axon terminals onto song premotor neurons in RA, allowing for further context-dependent regulation of the CBG pathway’s influence on song variability ^37,41^. More recently, we showed that pharmacologic blockade of alpha-adrenergic receptors in the CBG increases pitch variability of directed songs to the degree of undirected singing, while agonism of the same receptors results in reduced pitch variability during undirected singing. Intriguingly, in our more recent study, selective pharmacologic blockade of D1 receptors did not alter variability in either context, potentially due to the different time course of drug application (days vs hours). While additional experiments will be needed to better understand how DA and NA act in the CBG to influence song variability, it is clear that catecholaminergic signaling in the CBG pathway is important to how song is modulated to facilitate courtship.

### Hypothesis for opposing roles of NA and DA in context-dependent singing

Given the interactions between the dopaminergic and noradrenergic systems described here, a remaining parsimonious explanation is that NA and DA act in opposing manners to drive context-dependent differences in neural activity and behavior. For example, in the presence of a female, increased LC activity could result in release of NA in VTA/SNc, which could suppress DA neuron activity directly via α_2_ adrenergic receptors, or indirectly through glutamate release onto interneurons. This would be in concordance with previous observations made in VTA/SNc showing that interneurons, but not dopaminergic neurons, display increased IEG expression after directed singing. Such a reduction in VTA/SNc activity could then lead to decreased excitability in downstream areas such as Area X or LMAN. This suppression would be augmented by the direct effect of NA release onto the CBG, as indicated by the inhibitory effect of NA on Area X MSNs ^9^. Indeed, opposite effects of NA and DA have been reported before in mouse MSNs ^42^. Thus, based on our results we propose the hypothesis that such oppositional neuromodulatory systems may drive context-driven behaviors via CBG circuitry. Testing these ideas will ideally require improved viral genetic control over the NA and DA systems and will yield important advances to our understanding on how neuromodulatory systems interact to enable flexible complex behaviors such as learned birdsongs.

